## Supplementary Information for "A host recognition module shared among distant *Alteromonas* bacteriophage families features tail fibers with transient chaperone “caps”"

**Table S1.** Confidence scores of AlphaFold-generated models.

**Table S2.** Oligonucleotides used to generate protein expression plasmids.

**Table S3.** Percent similarity in the nucleotide sequence alignments of the proteins located in the host recognition module of phages V22, A5 and P24.

**Figure S1.** Genome comparative analysis of several N4-like schitoviruses that share a giant RNA polymerase of ca. 10 kb in size.

**Figure S2.** AlphaFold-based analysis of additional phage components: the central fiber, gp41 “Dit” of P24, short tail fiber gp23 of V22, CheY-like gp11 of A5, and the *Salmonella* phage S16 tail fiber IMC.

**Figure S3.** Sequence and structure alignment of the knob and head domains of the indicated tail fibers.

| Phage | Protein | Program used | pLDDT (average) | Interface pTM | Figure |
|---|---|---|---|---|---|
| A5 | gp8 (tail fiber) | Alphafold-Multimer | 75.8 | 0.57 | 4 |
| A5 | gp9 (chaperone) | AlphaFold 2.0 | 94.8 | n/a | 4 |
| A5 | gp9 (chaperone) | Alphafold-Multimer | 93.0 | 0.60 | 6 |
| A5 | gp8 Gly150 to Phe462 + gp9 | Alphafold-Multimer | 70.5 | 0.59 | 6* |
| V22 | gp26 (tail fiber) | Alphafold-Multimer | 77.4 | 0.67 | 4 |
| V22 | gp27 (chaperone) | AlphaFold 2.0 | 95.2 | n/a | 4 |
| V22 | gp27 (chaperone) | Alphafold-Multimer | 84.8 | 0.21 | 6 |
| V22 | gp26 Gly160 to Phe369 + gp27 | Alphafold-Multimer | 55.1 | 0.39 | 6* |
| P24 | gp44 (tail fiber) | Alphafold-Multimer | 75.1 | 0.65 | 4 |
| P24 | gp45 (chaperone) | AlphaFold 2.0 | 94.5 | n/a | 4 |
| GOV_bin2917 | gp51 (tail fiber) | Alphafold-Multimer | 65.5 | 0.46 | 4 |
| GOV_bin2917 | gp50 (chaperone) | AlphaFold 2.0 | 95.3 | n/a | 4 |
| A5 | gp6 (central fiber) | Alphafold-Multimer | 81.5 | 0.54 | S2 |
| A5 | gp11 (CheY-like) | AlphaFold 2.0 | 92.4 | n/a | S2 |
| V22 | gp24 (central fiber) | Alphafold-Multimer | 76.6 | 0.59 | S2 |
| V22 | gp23 (short tail fiber) | Alphafold-Multimer | 90.8 | 0.68 | S2 |
| P24 | gp43 (central fiber) | Alphafold-Multimer | 67.3 | 0.43 | S2 |
| P24 | gp41 (Dit) | Alphafold-Multimer | 76.1 | 0.51 | S2 |
| S16 | gp37 Ala516 to Lys749 (tail fiber) | Alphafold-Multimer | 94.3 | 0.92 | S2 |

**Table S1. Confidence scores of AlphaFold-generated models.** Confidence per residue is calculated as a predicted Local Distance Difference Test score (0-100), with an average of all residues within the models provided below. A pLDDT ≥ 90 have very high model confidence, residues with 90 > pLDDT ≥ 70 are classified as confident, while residues with 70 > pLDDT > 50 have low confidence. Interface pTM scores ("iptm+ptm") are a measure of predicted structure accuracy generated by AlphaFold-Multimer and provide the overall confidence score for the complete model (scored 0 to 1). *Models kindly generated by Petr G. Leiman (The University of Texas Medical Branch at Galveston, USA).

| Construct | Forward primer (5´- 3´) | Reverse primer (5´- 3´) |
|---|---|---|
| *pQE30_HGT* backbone | GCCCTGGAAATACAGATTCTCG | AATTAGCTGAGCTTGGACTCCT |
| *gp8_gp9* insert | CGAGAATCTGTATTTCCAGGGCATGGCTAGTACATTTTGGATT | AGGAGTCCAAGCTCAGCTAATTTTACCAAGTTAAGTTTGCTAGATAT |
| *gp8* insert | CGAGAATCTGTATTTCCAGGGCATGGCTAGTACATTTTGGATT | AGGAGTCCAAGCTCAGCTAATTTTAGAAGTTTACAGTGATTTTTGC |

**Table S2. Oligonucleotides used to generate protein expression plasmids.**

| | | V22 | | | | | | | | | | | A5 | | | | |
|---|---|---|---|---|---|---|---|---|---|---|---|---|---|---|---|---|---|
| | | gp21 | gp22 | gp23 | gp24 | gp25 | gp26 | gp27 | gp29 | gp30 | gp31 | gp32 | gp4 | gp5 | gp6 | gp8 | gp9 |
| A5 | gp4 | --- | n.s.s. | --- | --- | --- | --- | --- | --- | --- | --- | --- | --- | --- | --- | --- | --- |
| | gp5 | --- | --- | n.s.s. | --- | --- | --- | --- | --- | --- | --- | --- | --- | --- | --- | --- | --- |
| | gp6 | --- | --- | --- | 69.05%<br>Q.c. = 11% | --- | --- | --- | --- | --- | --- | --- | --- | --- | --- | --- | --- |
| | gp7 | --- | --- | --- | --- | 67.65%<br>Q.c. = 82% | --- | --- | --- | --- | --- | --- | --- | --- | --- | --- | --- |
| | gp8 | --- | --- | --- | --- | --- | 58.70%<br>Q.c. = 35% | --- | --- | --- | --- | --- | --- | --- | --- | --- | --- |
| | gp9 | --- | --- | --- | --- | --- | --- | 80.70%<br>Q.c. = 100% | --- | --- | --- | --- | --- | --- | --- | --- | --- |
| | gp10 | --- | --- | --- | --- | --- | --- | --- | 50%<br>Q.c. = 35% | --- | --- | --- | --- | --- | --- | --- | --- |
| | gp11 | --- | --- | --- | --- | --- | --- | --- | --- | 42.57%<br>Q.c. = 86% | --- | --- | --- | --- | --- | --- | --- |
| | gp12 | --- | --- | --- | --- | --- | --- | --- | --- | --- | 57.14%<br>Q.c. = 10% | --- | --- | --- | --- | --- | --- |
| | gp13 | --- | --- | --- | --- | --- | --- | --- | --- | --- | --- | 42.31%<br>Q.c. = 89% | --- | --- | --- | --- | --- |
| P24 | gp40 | n.s.s. | --- | --- | --- | --- | --- | --- | --- | --- | --- | --- | --- | --- | --- | --- | --- |
| | gp41 | --- | n.s.s. | --- | --- | --- | --- | --- | --- | --- | --- | --- | n.s.s. | --- | --- | | --- |
| | gp42 | --- | --- | n.s.s. | --- | --- | --- | --- | --- | --- | --- | --- | --- | n.s.s | --- | | --- |
| | gp43 | --- | --- | --- | 51.72%<br>Q.c. = 2% | --- | --- | --- | --- | --- | --- | --- | --- | --- | 50.91%<br>Q.c. = 68% | --- | --- |
| | gp44 | --- | --- | --- | --- | --- | 61.54%<br>Q.c. = 12% | --- | --- | --- | --- | --- | --- | --- | --- | 42.86%<br>Q.c. = 52% | --- |
| | gp45 | --- | --- | --- | --- | --- | --- | 41.86%<br>Q.c. = 21% | --- | --- | --- | --- | --- | --- | --- | --- | n.s.s. |

**Table S3. Percent similarity in the nucleotide sequence alignments of the proteins located in the host recognition module of phages V22, A5, and P24.** Comparisons were done using the tBLASTx suite. Q.c.: Query cover.

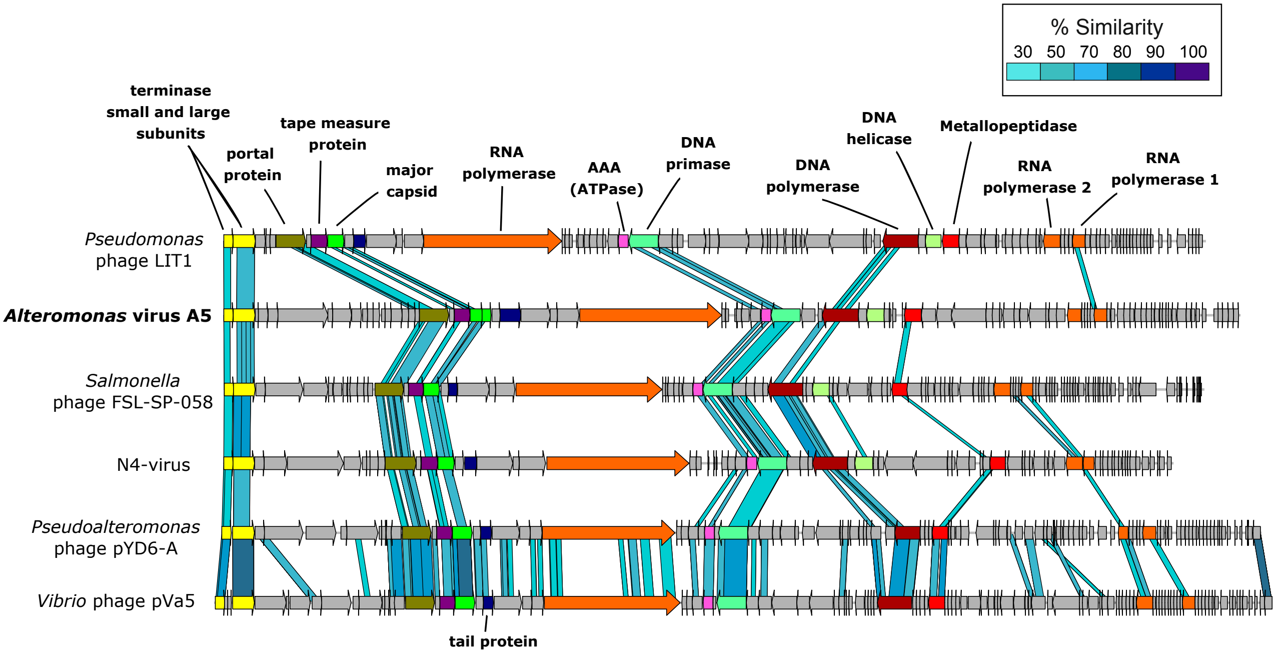


**Figure S1. Genome comparative analysis of several N4-like schitoviruses that share a giant RNA polymerase of ca. 10 kb in size.** Sequence comparisons were performed using tBLASTx with 30% minimal similarity on 100 bp minimum alignments. The host-recognition module of these phages did not show synteny or sequence similarity.

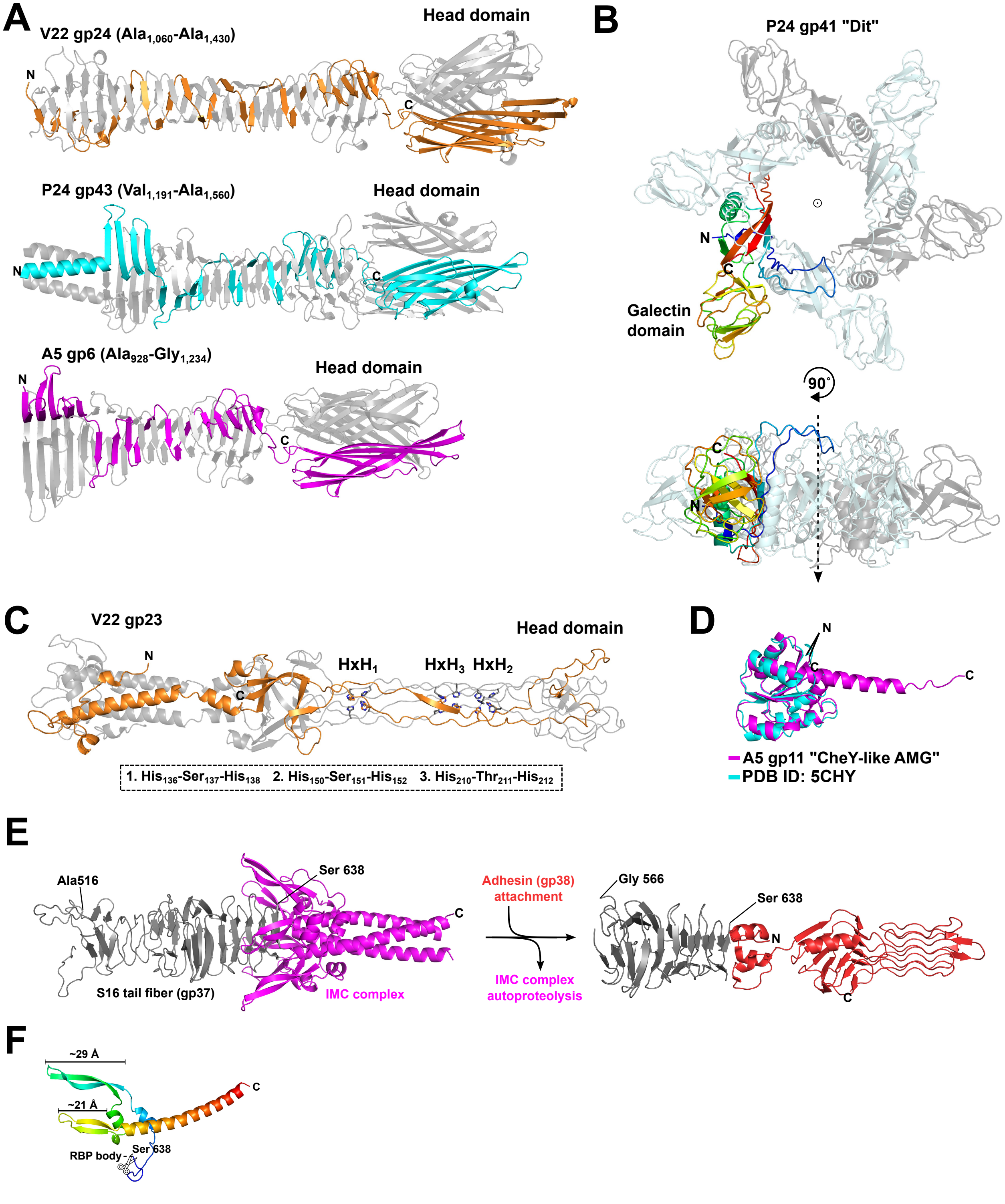


**Figure S2.** See next page

**Figure S2. AlphaFold-based analysis of additional phage components: the central fiber, gp41 "Dit" of P24, short tail fiber gp23 of V22, CheY-like gp11 of A5, and the *Salmonella* phage S16 tail fiber IMC.** **(A)** Ribbon diagrams of the homotrimeric C-terminal tips of the putative central RBP of phages V22 (gp24), P24 (gp43), and A5 (gp6) predicted using AlphaFold-Multimer. The distal head domains of all three central RBPs consists of extended lectin-like binding domains similar to the head domains of the phage's tail fibers (e.g., A5 gp8). **(B)** Ribbon diagram of the hexameric assembly of P24 gp41 predicted using AlphaFold-Multimer which resembles a classical "evolved" distal tail protein (Dit) complex, which is a highly conserved central building block of all Siphoviral baseplates. As highlighted, gp41 features a galectin-like carbohydrate-binding module (CBM) which is a characteristic feature of "evolved" Dit complexes. **(C)** Ribbon diagram of homotrimeric V22 gp23 as predicted using AlphaFold-Multimer features a classic phage T4 gp37-like tail fiber architecture (PDB ID: 2XGF) with an intertwined, elongated needle-like domain containing three HxH metal-binding sites that are expected to each be occupied by an $Fe^{2+}$ ion. **(D)** Ribbon representations of A5 gp11 and *E.* coli chemotaxis protein CheY (PDB ID: 5CHY) superpose with a root-mean-square deviation (C-α RMSD) of 1.67 Å. **(E)** Ribbon diagram of the *Salmonella* phage S16 tail fiber (gp37) and its autoproteolytic C-terminal IMC predicted using AlphaFold-Multimer. The fiber tip associates with its adhesin (gp38) to form a mature, functional fiber (to the right, PDB ID: 6F45 [45]. **(F)** The gp37 IMC closely resembles the IMCs of the T5 and K1F RBPs (shown in **Fig. 6 A,C**), but was predicted as having an additional β-hairpin domain. All model confidence scores are provided in **Table S1**.

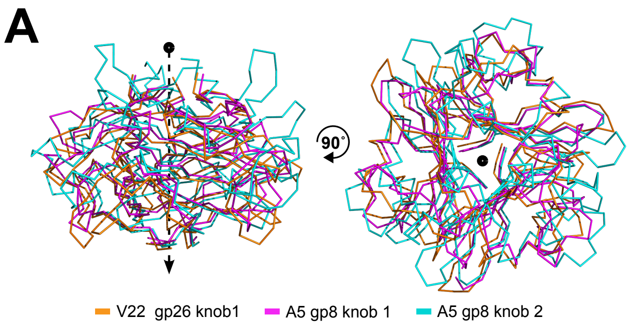


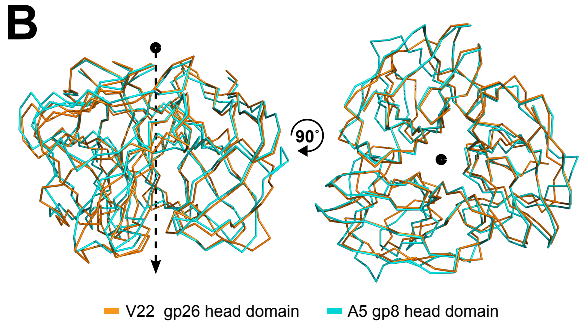


**C**

```
                   1        10        20        30        40        50        60        70        80        90        100      109
                   |--------+---------+---------+---------+---------+---------+---------+---------+---------+---------+--------|
               A5_        AFGFGGISPAQSPNGDLNDIERTGVYDVDGATLNTPPAFNPKNGMCLHLSRTDTLATQLLYSRRTTQMFMRVNEGSGWENWAEFYH
GOV_bin2917_Knob1         AFGLGGAVIENF--ADFNDISETSFIGGNNTILNAPPWMDSRVMGGIHVDKGGR--TLQIMPSRFGDLGFRTSFDNVYGDWQEIYH
         A5_Knob2     VNPLDYGIGIHRHNDGSPTEMKNVSADALAGGLYTGYEGLN--NNATLGDNPFTDSGAAYGLLNITGVDPDPTKYIFQLAGQYSGYRLRWRAATVSGFSTWRDVYH
GOV_bin2917_Knob2     VNPLNYGIKTHA-NLPSISDLNTIGASLLTSVVDSSTQNRP--DSAVGHVINLSQNNAANSQIVTDRIYNDIWFRTSSGSVYKDFSKLYH
        P24_Knob1  AWGTGLLVGGAIKNS-EYSVVDDLNESSQPSQFFVFDSATLNRP--STPEGAVI---PNGVGISLSRTSNIWGQLTLSRAGGSVGTLITRVKDGTTVVERLAYD
        V22_Knob1         AFGVGVNQTTIISDINAVTDGSSGFYGINNGTIGKPEFGGNTFGVVTKFPYTNF------------QPQLFTGSVGTGNPTAAIRVYDDLTSSFSDWLEFHH
        Consensus  .......afG.g........s..#..........g........p....n..g.v..f.................#p...t.............r......s.fs.w....h
```

| | V22_Knob1 | A5_Knob2 | GOV_Bin2917_Knob1 | A5_Knob1 | GOV_Bin2917_Knob2 | P24_Knob1 |
|---|---|---|---|---|---|---|
| V22_Knob1 | 100 | 17.24 | 20.51 | 19.51 | 18.99 | 17.72 |
| A5_Knob2 | 17.24 | 100 | 21.52 | 16.87 | 18.39 | 14.29 |
| GOV_Bin2917_Knob1 | 20.51 | 21.52 | 100 | 27.16 | 23.68 | 21.79 |
| A5_Knob1 | 19.51 | 16.87 | 27.16 | 100 | 26.25 | 23.75 |
| GOV_Bin2917_Knob2 | 18.99 | 18.39 | 23.68 | 26.25 | 100 | 25 |
| P24_Knob1 | 17.72 | 14.29 | 21.79 | 23.75 | 25 | 100 |

**Average Seq. Similarity = 20.8 % ± 2.8**

**D**

```
                  1        10        20        30        40        50        60        70        80        90        100   106
                  |--------+---------+---------+---------+---------+---------+---------+---------+---------+---------+-----|
          A5_Head NFNEFGG-NGNECIATGEIVTSTIARFYLPINAKTKPVSLSIIGSFNVFRLSGSVVSGAGTGSFALSAISSPKMAVIDVTGS-GLTLGEVVNLRCAVNNAKITVNF
 GOV_Bin2917_Head NFNEFGGANAGDVIALGESVTSTLARFYLPINGVIQPSSLSLIGSVSVFK--AAVALESNVTNVGLSGISGRKVAVVDVTCSVGLTVGDQLNLRMPTSSDKITINF
         V22_Head NFNEFGG-NVGDTIATGAAHGANYASFELPINSTSSPAGVSVAGAFSVVAKSGTATRVASTTVNFSGADSSKKIAVVSVTSSGAFTAGEVLILRSAANNSKITVNF
         P24_Head NFNEFGGAAGDNIGCIGFAVSTTVADFYFPLVSKAPPTGRGVVGTFDVVTPDGTLI-AAGVTPSFNGRSSFREL-VLQVSGLSGITKDQPLKLRCKTATDKITAQF
        Consensus NFNEFGGang.#.ia.G.av..t.A.FylPinsk..P.g.sv.G.f.Vv...gt....agvt..f.g..S.rk.aV..Vtgs.g.T.g#.l.LRc.t..dKIT.#F
```

| | P24_Head | GOV_Bin2917_Head | A5_Head | V22_Head |
|---|---|---|---|---|
| P24_Head | 100 | 33.33 | 36.27 | 33.98 |
| GOV_Bin2917_Head | 33.33 | 100 | 55.88 | 41.75 |
| A5_Head | 36.27 | 55.88 | 100 | 49.04 |
| V22_Head | 33.98 | 41.75 | 49.04 | 100 |

**Average Seq. Identity = 41.7 % ± 8.3**

**Figure S3. Sequence and structure alignment of the knob and head domains of the indicated tail fibers. (A)** Superposition of the knob domain from V22 gp26 (orange) with the two knob domains of A5 gp8 (knob 1, magenta; knob 2, cyan) was performed using the DALI server [38]. Despite all knob domains sharing low (<20%) sequence identity, they feature the same domain structure with root-mean-square deviations (RMSD) of 1.5 Å and 2.3 Å for the V22 knob to knob 1 and knob 2 of A5, respectively. **(B)** Superposition of the distal head domains from V22 gp26 (orange) and A5 gp8 (cyan) also revealed they share the same domain structure with RMSD of 0.59 Å as was expected from 49% sequence similarity. MultAlin-generated [74] sequence alignments of all knob domains **(C)** and all distal head domains **(D)** from the four phage TFs described in this study with average sequence similarities determined using Clustal Omega [81]. All model confidence scores are provided in **Table S1**.
